# KGBN: Augmenting and optimizing logical gene regulatory networks using knowledge graphs

**DOI:** 10.64898/2026.01.29.702644

**Authors:** Luna Xingyu Li, Yue Zhang, Boris Aguilar, Tazein Shah, John H Gennari, Guangrong Qin

## Abstract

Logical gene regulatory network (GRN) models provide interpretable, mechanistic representations of cellular regulation and are widely used in systems biology. However, most existing models remain incomplete, context-specific, and difficult to extend to comprehensive GRNs, limiting their broader applicability to tasks such as drug-response prediction and precision medicine. Here, add some ideas about virtual twins: With AML patients, we can use comprehensive GRNs to predict drug response. We present KGBN (Knowledge Graph–augmented Boolean Network modeling), a computational workflow for systematically augmenting logical GRN models. KGBN incorporates regulatory interactions derived from curated knowledge graphs as alternative logical rules while preserving the validated structure of existing models. Rule probabilities are optimized against experimental data to represent regulatory uncertainty and achieve data-driven calibration. Applying KGBN to acute myeloid leukemia, we show that extending an existing GRN with drug-target pathways and training against *ex vivo* drug-response data yields mutation-specific models that recapitulate known therapeutic sensitivities and signaling dependencies, demonstrating the utility of KGBN for interpretable, context-aware GRN modeling. (Should be predict)

## Introduction

Systems biology aims to explain how molecular components interact to produce emergent cellular behaviors, disease phenotypes, and treatment responses. Mathematical models play a central role in this effort by translating biological knowledge into executable representations that can be simulated and tested (1; 2). Among them, logical models have become a widely adopted formalism for representing gene regulatory networks (GRNs) by describing the qualitative activation states of genes and proteins and using logical rules to capture their regulatory relationships (3–5).

GRN modeling has been instrumental in revealing the mechanisms of complex diseases. In acute myeloid leukemia (AML), for example, logical models have been used to capture hematopoietic stem cell differentiation and the dysregulation of transcriptional programs driven by oncogenic mutations such as *FLT3, NPM1*, and *RUNX1* (6–8). They have been used to study a wide range of biological processes, including signaling crosstalk, drug response, and resistance mechanisms in various diseases (e.g., breast cancer (9) and pancreatic cancer (10)). However, these models are typically constructed for specific biological questions, often omitting drug targets or signaling intermediates, and encode a single deterministic regulatory logic per node (11; 12). These limitations hinder their ability to represent regulatory uncertainty, capture patient heterogeneity, or adapt to new experimental contexts such as drug perturbation screens.

Recent advances provide an opportunity to address these challenges. First, biomedical knowledge graphs systematically curate causal relationships among genes, proteins, drugs, and phenotypes, offering a scalable resource for identifying missing regulatory interactions and drug–target connections (13; 14). Second, probabilistic Boolean networks (PBNs) extend classical Boolean models by allowing multiple alternative regulatory rules per node, thereby explicitly representing uncertainty, context dependence, and competing biological hypotheses (15; 16). Third, large-scale experimental datasets, including ex vivo drug-response and multi-omics profiling, enable data-driven calibration of model parameters, transforming static logic models into quantitatively informed predictive systems.

Here, we introduce KGBN (Knowledge Graph-augmented Boolean Network modeling), a computational framework that integrates these elements into a unified workflow for augmentation and optimization of GRN models. KGBN uses knowledge graphs to augment existing logical models with disease- and drug-relevant regulators, encodes alternative regulatory hypotheses using PBNs, and optimizes rule probabilities against experimental data to improve predictive fidelity. We demonstrate the utility of this framework through drug response prediction in AML and through reproduction of a published pancreatic cancer model, illustrating both its predictive capability and its reliability as a general execution and extension platform for GRN models. With the potential to apply to diverse biological and disease contexts, this framework will facilitate mechanistic hypothesis generation, mutation-specific drug response analysis, and *in silico* exploration of therapeutic combinations.

## Results

### The KGBN toolkit

We developed KGBN, an open-source Python library for Boolean network and probabilistic Boolean network modeling, extension, optimization, and analysis, available at https://github.com/IlyaLab/KGBN. It supports the full workflow summarized in Figure 1 and described in the Methods. Its core functionality includes:

1. Network construction and manipulation: Load or build models from SBML-qual, text, or connectivity matrices; merge and extend networks;
2. Boolean network modeling: Perform deterministic or stochastic updates, steady-state analysis, and trajectory simulation;
3. Probabilistic modeling: Simulate PBNs stochastically, perform steady-state analysis, and visualize the network;
4. Knowledge-graph integration: Query KGs to derive models from causal relations between biochemical entities and phenotypes;
5. Optimization and evaluation: Fit model predictions to quantitative experimental data, and evaluate on test cases.

**Fig. 1.**
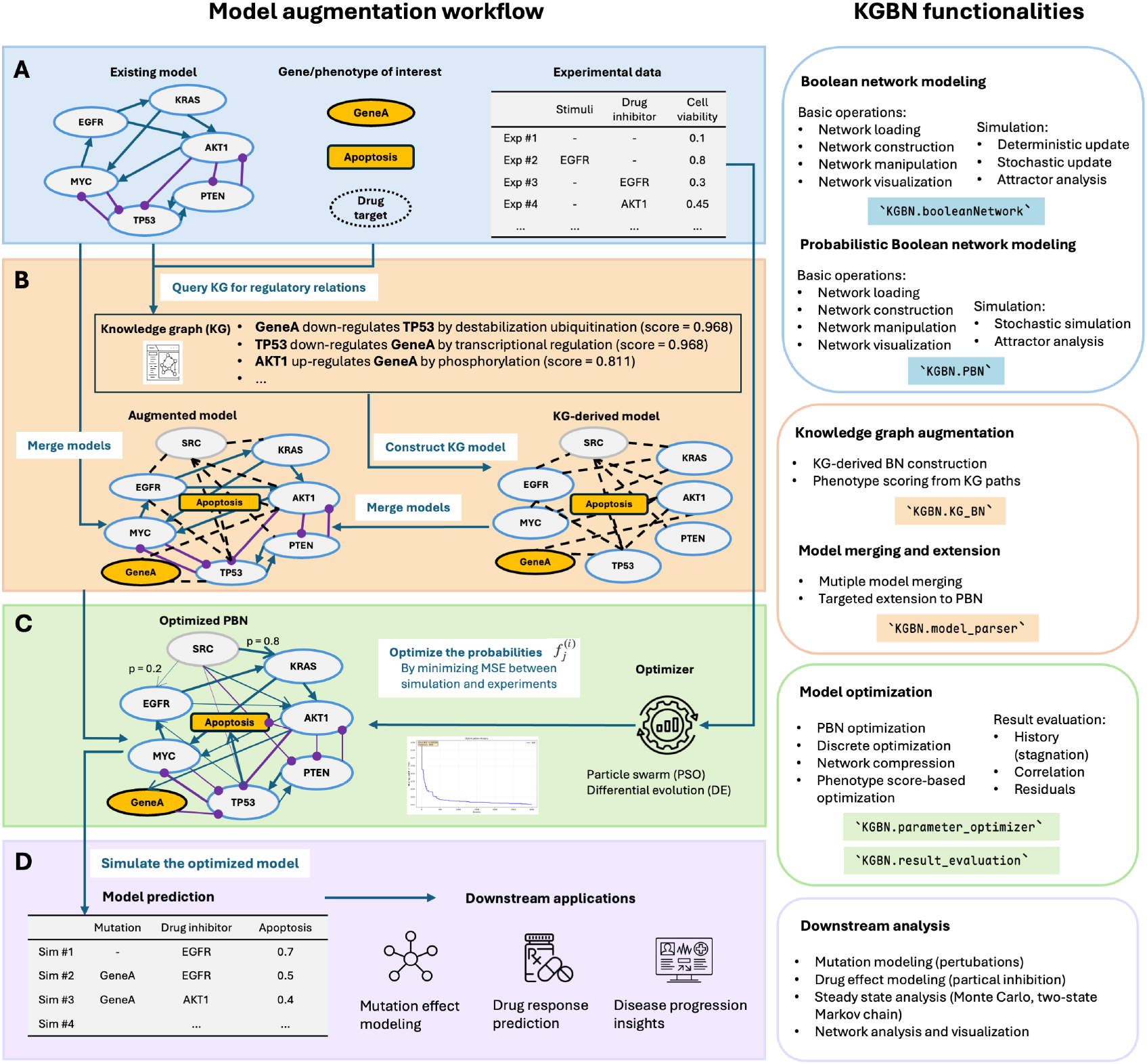
Overview of the workflow and KGBN core functionalities. **a** Identifying and standardizing existing models, genes, phenotypes, and experimental data. **b** Extending the model via knowledge graph (KG) and probabilistic Boolean networks (PBNs). **c** Optimizing the PBN using experimental data. **d** Simulating and evaluating the optimized model.

To demonstrate the correctness and practical applicability of the toolkit, we evaluated KGBN on a published logical model of pancreatic cancer signaling (17). Using the same model structure and simulation protocol reported in that study, we reproduced the published simulation-based ranking of drug combinations (Supplementary Fig. S1). This reproduction provides a validation benchmark for the software and a foundation for the AML use case below.

### A use case: AML drug response prediction

We next applied KGBN to a published Boolean model of AML to test whether knowledge-guided augmentation and probabilistic rule selection can recover drug-response phenotypes across mutational contexts (7). The starting model captures major AML regulators but lacks parts of the signaling circuitry needed to represent several targeted therapies. KGBN therefore augments the network with KG-supported interactions and converts affected nodes into PBN junctions whose rule probabilities are learned from drug-response data.

#### Knowledge-guided augmentation predicts drug response

To test whether one shared regulatory logic could explain responses to multiple therapies, we optimized a single augmented PBN against the combined response profiles of five targeted AML drugs. As summarized in Figure 2A,B, the KG-augmented network adds missing drug-target circuitry and increases connectivity around major survival regulators, most visibly through the introduction of *BTK* and *SYK* and their links to nodes such as *MAPK1, AKT1*, and *MYC*. These additions place the original AML model in a more realistic therapeutic context, where FLT3-centered signaling converges with MAPK, AKT, and STAT5 survival programs and where SYK and BTK associated pathways can modulate drug response (18–20).

**Fig. 2.**
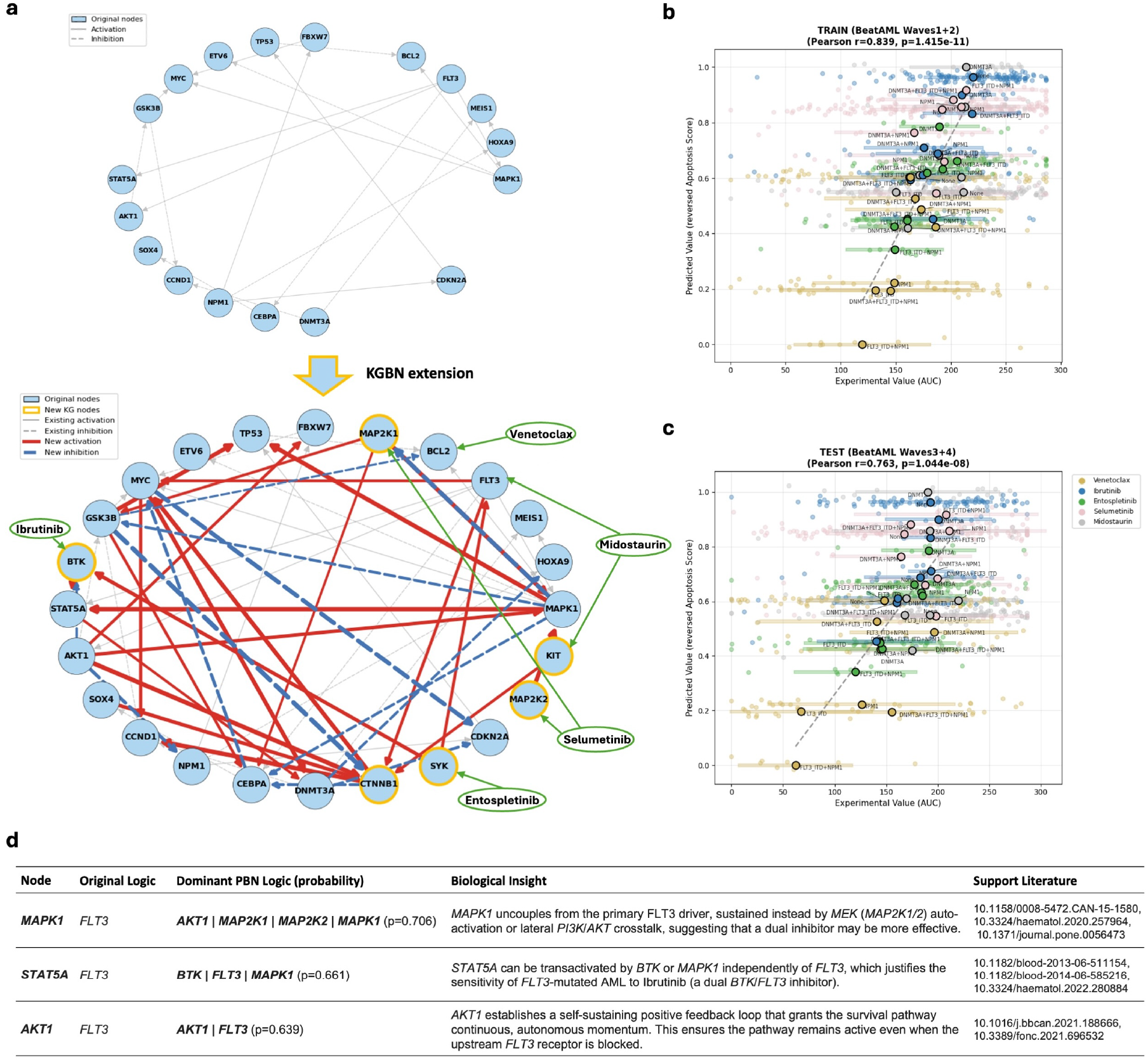
Knowledge-guided augmentation and optimization of an AML model for drug-response prediction. **a** Augmention of the AML Boolean network, with newly introduced nodes and edges highlighted. **b, c** Correlation between measured Beat AML AUC and the predicted reversed apoptosis score in the training cohort (Waves 1+2) and test cohort (Waves 3+4), respectively. Each dot represents a mutation-defined patient group and is colored by drug. **d** Comparison of representative logical rules in the original and optimized models, highlighting rewiring of *MAPK1, STAT5A*, and *AKT1*.

After training on drug response seperated by mutation status of *FLT3*-ITD, *NPM1* and *DNMT3A*, the augmented model remained aligned with measured drug response in the Beat AML training cohort, in the held-out Beat AML test cohort, and after transfer to the independent FPMTB dataset (Figure 2C–E; panel statistics shown in the figure). The main mechanistic insight comes from Figure 2F, which compares the dominant rules for three key signaling hubs before and after optimization. *MAPK1* shifts away from strict *FLT3* dependence toward a permissive rule that can be sustained through *AKT1, MAP2K1, MAP2K2*, and self-maintenance, consistent with effector uncoupling and rapid bypass of receptor-level inhibition. *STAT5A* similarly moves from exclusive *FLT3* control to a redundant *BTK*/*FLT3*/*MAPK1* gate, providing a mechanistic explanation for why BTK-directed therapy can suppress a survival axis that would be invisible in a purely FLT3-centered model. Finally, *AKT1* acquires self-sustaining logic, consistent with the persistence of AKT signaling through positive-feedback circuits that continue to support leukemic survival after upstream blockade. Together, these rule shifts point to a resistant network architecture in which AML cells preserve viability by distributing control across parallel MAPK, BTK/STAT5, and AKT modules rather than relying on a single dominant receptor input (20; 21).

#### LOCO validation reveals generalizable logic

We next asked whether the optimized rule probabilities reflect generalizable regulatory logic rather than cohort-specific noise by performing leave-one-mutation-out (LOCO) cross-validation across mutation-defined groups. Predictions for mutation profiles that were completely withheld during optimization remained informative overall (**??**A–C), with the best prediction observed for isolated *DNMT3A* and *NPM1* states. This pattern is biologically consistent with the idea that single-mutation contexts can be represented as modular perturbations of a shared signaling backbone, whereas the wild-type baseline and more complex multi-mutant states depend more strongly on cooperative rewiring that is harder to infer without direct contextual data.

The rule-probability heatmap in **??**D reinforces this interpretation. Some nodes preserve a stable core logic across folds, such as the near-complete retention of the original rule for *NPM1*, whereas integrative nodes such as *MAP2K1* repeatedly favor KG-derived inhibitor-wins logic. The model consistently recovers a mixture of conserved regulatory backbone and context-dependent escape logic. It is expected if the optimized PBN were capturing biologically reproducible pathway structure rather than arbitrary dataset-specific fits.

#### NRAS extension reveals Ras-driven rewiring

Finally, we asked whether KGBN can be further leveraged to augment the model with an additional recurrent mutation that is well known to reshape therapeutic response. As a proof of concept, we extended the AML network with *NRAS* and the intermediary regulator *DAB2IP* (Figure 3A,B), creating explicit routes by which oncogenic Ras signaling can connect to *AKT1, MAPK1, GSK3B*, and *TP53*. By training the extended model on drug response grouped by *FLT3*-ITD, *NPM1, DNMT3A*, and *NRAS* mutation status, we obtained an optimized model that covers additional patient mutation profile (purple-colored nodes in **??**C-E), allowing direct representation of a clinically relevant relapse-associated bypass program.

**Fig. 3.**
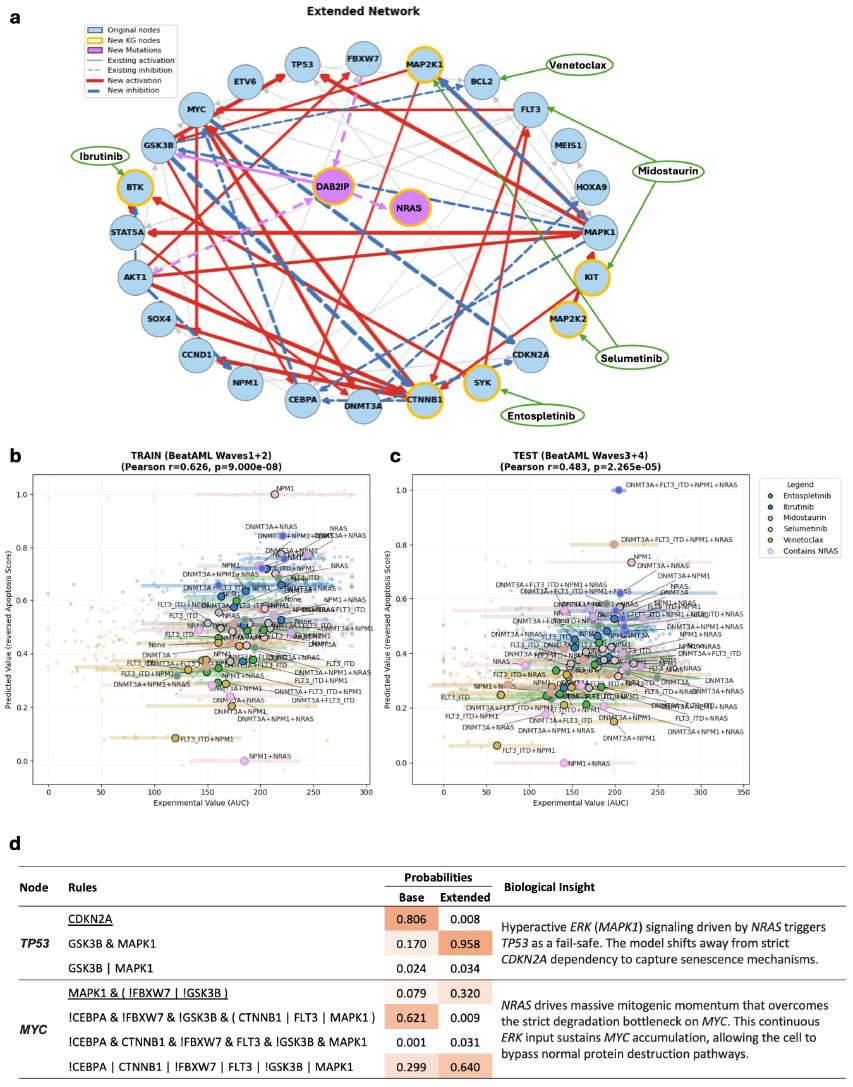
Extension of the AML model with *NRAS*. **a** Network extended by adding *NRAS, DAB2IP*, and their KG-supported interactions. **b**–**c** Correlation between measured drug response and predicted apoptosis score for the NRAS-extended model across the training cohort and test cohort. **d** Comparison of representative rule probabilities in the NRAS-extended model and the previous model, highlighting rewiring of *TP53* and *MYC*.

We compared the two PBNs before and after *NRAS*-extension in Figure 3F ((Supplementary Table. S2)). In the baseline model, *TP53* is controlled primarily through *CDKN2A*/ARF-associated stabilization. After *NRAS* augmentation, *TP53* becomes dominated instead by a *GSK3B*-*MAPK1* rule, consistent with oncogene-induced stress signaling in which excessive Ras/MAPK flux is interpreted as DNA damage or replication stress and is routed directly into senescence or apoptosis associated control. At the same time, *MYC* shifts from a stringent rule that required simultaneous escape from several degradation constraints to a far more permissive logic. This rewiring is consistent with sustained Ras/MAPK activity stabilizing MYC and allowing proliferative drive to persist even when the normal *GSK3B*/*FBXW7*-dependent degradation machinery is not fully disabled. Together, these two changes show that *NRAS* does not simply add another proliferative branch to the network; it reorganizes both the stress-response module (*TP53*) and the proliferative module (*MYC*), creating a more Ras/MAPK-dominated state with reduced dependence on upstream receptor control (21–24).

This interpretation also helps explain why *NRAS*-mutant AML can be especially difficult to suppress with therapies aimed at a single upstream target. By wiring Ras signaling through *DAB2IP* into both survival and stress-response circuits, the extended model suggests that mutation acquisition produces network-wide rewiring rather than a simple additive effect. In that setting, restoring durable drug sensitivity is likely to require interrupting the downstream Ras/MAPK and AKT escape routes that become dominant after the mutation is introduced.

## Discussion

In this study, we introduced KGBN, a knowledge graph augmented framework to improve logical GRN models using probabilistic representations and experimental calibration. Applying the framework to AML, we demonstrated that augmenting an existing GRN with drug-target pathways and optimizing rule probabilities against *ex vivo* drug-response data allows the model to capture mutation-specific therapeutic sensitivities and decipher signaling dependencies.

A key strength of KGBN lies in its explicit representation of regulatory uncertainty. Rather than committing to a single deterministic logic, the PBN formulation allows multiple competing regulatory hypotheses to coexist, with their relative influence learned from data. In the AML use case, the optimized rule probabilities were reproducible across cohort splits and LOCO folds, and sensitivity analysis highlighted a compact set of control points that dominate the phenotype prediction. This illustrates how data-trained PBNs can serve not only as predictive models but also as interpretable summaries of pathway utilization and parameter identifiability. Further, by grounding newly introduced nodes and edges in curated knowledge resources, KGBN provides transparent and traceable biological provenance for model augmentation. This is particularly valuable for modeling drug mechanisms of action and resistance, where regulatory context and pathway crosstalk play a critical role. Moreover, the phenotype scoring strategy enables alignment between qualitative network states and quantitative experimental readouts, bridging a longstanding gap between logical modeling and experimental data.

KGBN complements our prior work on LM-Merger, which integrates multiple existing GRN models to improve robustness and regulatory coverage (25). While LM-Merger demonstrated improved generalization by combining AML and hematopoiesis models (7; 26), KGBN addresses a complementary setting in which curated knowledge graphs are used to extend a single validated model with new genes, pathways, and regulatory hypotheses. Together, these approaches support flexible GRN construction through either model–model integration or knowledge-driven extension, depending on data availability and modeling goals.

Although this study focuses on AML as a primary use case, the workflow is not disease-specific. We have demonstrated its generalizability by reproducing a published pancreatic cancer logical model. Building on this validated baseline, we are currently augmenting the pancreatic cancer model with drug-target pathways and probabilistic logic to enable data-driven prediction of combination therapies. Beyond extending existing models, KGBN can use knowledge graphs to extract regulatory topology directly from a set of genes and phenotypes of interest and construct novel logical models for system-level simulation. More broadly, the modular design of KGBN allows integration of additional structured knowledge sources beyond SIGNOR. As a future direction, we are actively incorporating causal and drug-response knowledge from NCATS Translator resources to expand regulatory coverage, improve drug-target representation, and support cross-disease modeling (13). These extensions will further strengthen the workflow’s applicability to diverse contexts and advance its role as a general framework for GRN modeling.

We would like to acknowledge a few limitations of our study. First, the quality of model extension depends on the completeness and accuracy of underlying knowledge graphs, which may contain conflicting or context-agnostic interactions. While the probabilistic framework partially mitigates this issue, our future work will incorporate evidence weighting, disease-specific filtering, or ensemble knowledge graph integration. Second, optimization against limited datasets risks overfitting, particularly for large networks. With the data we have, we have carried out cross-validation and external validation with another independent dataset. Another strategy to avoid overfitting would be to try model compression strategies, especially as models become larger.

Overall, KGBN advances logical GRN modeling by providing a reproducible, extensible, and data-informed pathway from curated mechanistic knowledge to predictive, context-aware models. By enabling systematic incorporation of drug targets, regulatory uncertainty, and experimental calibration, this framework lays the groundwork for scalable mechanistic modeling in systems biology and supports the broader vision of precision medicine.

## Methods

### Boolean networks and PBNs

We model a gene regulatory network as a discrete dynamical system composed of binary state variables. Let *V* = {*x*_1_, …, *x*_*n*_} denote the set of nodes, where each node *x*_*i*_(*t*) ∈ {0, 1} represents the activity state of a gene, protein, or phenotype at discrete time *t*. In a Boolean network, the state of each node is updated by a Boolean function *x*_*i*_(*t* + 1) = *f*_*i*_(*x*_1_(*t*), …, *x*_*n*_(*t*)), where *f*_*i*_: {0, 1}^*n*^ → {0, 1} encodes the regulatory logic of node *i*. The global network state is given by **x**(*t*) = (*x*_1_(*t*), …, *x*_*n*_(*t*)). In KGBN, we adopt a synchronous update scheme, where all node states are updated simultaneously at each discrete time step.

PBNs generalize this formulation by allowing multiple alternative update rules per node. For each node *x*_*i*_, we define a set of candidate Boolean functions 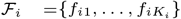 together with an associated probability vector 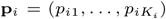, 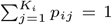. At each update step, one function *f*_*ij*_ is selected according to **p**_*i*_, and the network state is updated accordingly. This stochastic rule selection induces a Markov chain over the global state space {0, 1}^*n*^.

In KGBN, this formulation provides a natural mechanism for incorporating regulatory information derived from knowledge graphs. The original Boolean rule of a node is preserved as one element of ℱ_*i*_, while additional rules derived from knowledge-graph interactions are introduced as alternative elements of ℱ_*i*_. The probability vector **p**_*i*_ therefore controls the relative contribution of competing regulatory hypotheses.

The steady-state distribution ***π*** is defined as the invariant distribution of the Markov chain induced by the PBN, satisfying ***π*** = ***π****P*, where *P* is the state transition matrix determined by probabilistic rule selection (27). Steady states in KGBN are estimated using either Monte Carlo simulation, by sampling long stochastic trajectories, or a two-state Markov chain approximation for ergodic PBNs, enabling efficient evaluation of model outputs during rule-probability optimization (15).

### The KGBN workflow

KGBN is designed as a modular workflow that augments existing logical gene regulatory network models with knowledge-driven structure and data-driven calibration. As illustrated in Figure 1, the workflow proceeds from standardized model and data preparation (a), through systematic network augmentation using curated causal knowledge (b), to probabilistic model optimization (c) and downstream simulation (d). Users can adapt the workflow to different disease contexts, data types, and modeling objectives. The following subsections describe each step of the workflow in detail.

#### Identifying and standardizing models

The workflow starts with collecting existing models based on the biological questions. We identify additional genes and phenotypes of interest that were not included in the model, including frequent mutated genes, key regulators of disease, drug targets, and phenotypes linked to experimental observations.

Conventional names are often used in models, e.g., *RAS*, a gene family involved in cell growth and cancer development that includes *KRAS* and *NRAS*. To facilitate knowledge graph lookup, all entities in the network should be normalized to their standard formats. For genes, KGBN accepts either standard names/symbols in HGNC (for human models) and NCBI Gene (for other species), or gene IDs in NCBI; for proteins, it accepts their IDs or names as in UniProt.

#### Extending the model via knowledge graphs

To augment existing GRN models, we develop a pipeline to query the SIGNOR database (the SIGnaling Network Open Resource) (28) and construct Boolean network models from curated molecular interactions (Figure 4). Given a query list of genes or proteins, we identify their interactions by computing the Steiner subgraph (29). This minimal connected subgraph contains all query nodes and their shortest paths through the knowledge graph. Within the extracted subgraph, we classify each directed edge as activating or inhibiting. We then apply a configurable joiner function to generate Boolean update rules:

- OR: the node is active if at least one activator is active.
- AND: the node is active only if all activators are active.
- Inhibitor wins: if any inhibitor is active, the target is inactive regardless of the activator state.
- Majority(Plurality): if the number of active activators is greater than (or equal to) the number of active inhibitors, then the target is active.

**Fig. 4.**
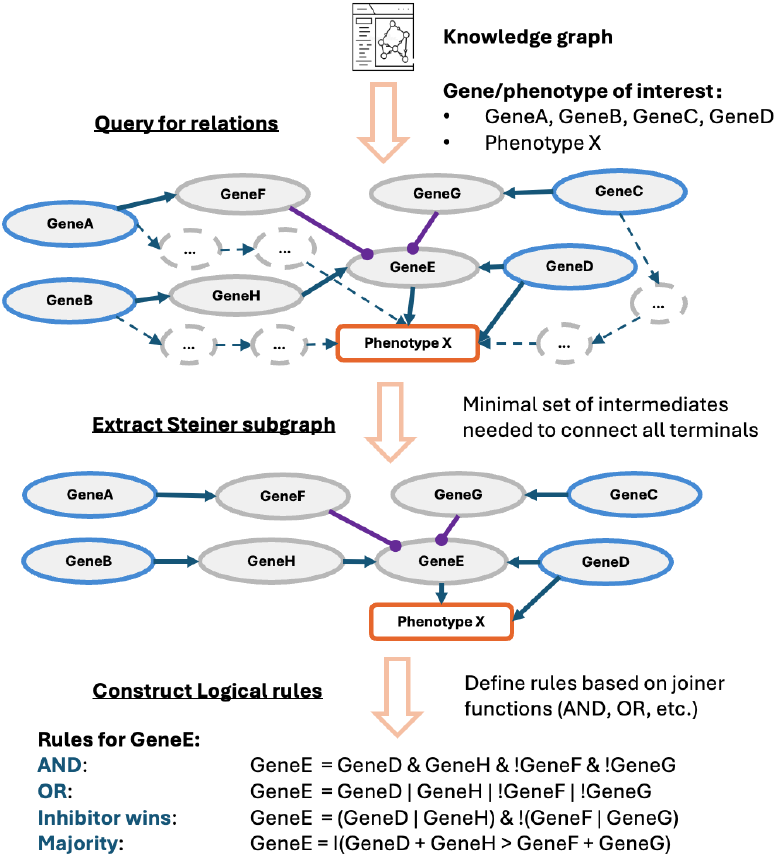
KG–based model extension in KGBN. Query genes (blue-colored) and phenotype (orange) of interest are connected via mediators (gray) based on their regulatory relations from the KG. Steiner subgraph extraction retains the minimal set of intermediate nodes and signed causal edges by removing extra ones (dashed). The resulting subgraph is converted into Boolean rules using configurable joiner functions. Example rules are shown for GeneE, the important intermediate node identified from the KG.

The knowledge graph query process includes both the genes present in the original model and additional genes of interest, allowing us to retrieve both the direct interactions among the new components and their connections to the pre-existing model. The resulting network captures a broader range of molecular events that influence phenotypes and enables simulations that more accurately reflect mutation–drug response relationships and uncover regulatory dependencies that may represent novel therapeutic targets.

To help quantify the impact of the network on phenotypes, we leverage the ProxPath framework (30). ProxPath quantifies how close genes are to a phenotype by filtering significantly short paths based on Z-score distribution of all relevant paths in SIGNOR. We take the resulting significant relationships and compute a phenotype score directly from simulated node states: regulators whose net path sign is positive are added, while regulators with a net negative sign are subtracted. This yields a quantitative and interpretable signed sum for each phenotype, ready for connecting network simulations with experimental data in the next step.

#### Probabilistic Boolean Networks extension

We consider two complementary strategies for incorporating new regulatory information derived from knowledge graphs into existing logical models. One strategy fully merges the base model with a knowledge graph–derived network, combining overlapping and unique nodes and regulatory rules into a single, expanded Boolean network. This approach, implemented in our previously described LM-Merger framework, produces a unified model with increased coverage and integrated logic across components (25). In this work, we focus on a more targeted extension strategy based on PBNs, where alternative regulatory rules derived from knowledge graphs are introduced, while the validated structure of the original model is still preserved.

Let the base model be a Boolean network defined on nodes *V* = *x*_1_, …, *x*_*n*_, where each node *x*_*i*_ is associated with a Boolean update function 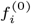. For each node *x*_*i*_ affected by model extension, we construct a set of alternative Boolean rules 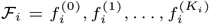, where 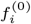 is the original rule from the base model and 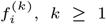, are additional rules derived from knowledge-graph interactions. Each rule set ℱ_*i*_ is associated with a probability vector 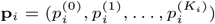 subject to the constraint 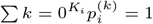.

This formulation yields a probabilistic Boolean network in which stochastic selection among alternative rules induces a Markov chain over the global state space {0, 1}^*n*^. The PBN explicitly represents regulatory uncertainty by allowing multiple competing mechanisms to coexist, with their relative influence controlled by the probability vectors **p**_*i*_. This structure provides a principled bridge between curated mechanistic knowledge and data-driven model calibration.

#### Optimizing the PBN using experimental data

To calibrate the extended PBN against experimental observations, we optimize the rule probability vectors {**p**_*i*_} using quantitative phenotype measurements. Each experimental condition *m* ∈ *C* is defined by a set of node perturbations (e.g., gene inhibition or activation), their efficacies, and an observed phenotype value *y*_*m*_, such as drug response or cell viability. The predicted output for condition *m* is denoted by *ŷ*_*m*_(**p**) and is obtained as a signed linear combination of steady-state node activation probabilities.

The optimization problem is formulated as minimization of the mean squared error between simulated and observed phenotype values across all experimental conditions:

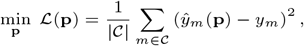

subject to 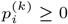 and 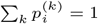 for all nodes *i*.

Because the objective is non-convex and the steady-state distribution depends on **p**, we employ global optimization strategies, specifically particle swarm optimization and differential evolution (31). Early stopping is applied when improvements in the objective fall below a predefined threshold. Model performance is assessed using prediction–observation correlations, residual analysis, and convergence diagnostics to ensure improved fidelity without overfitting.

#### Simulation and evaluation

After optimization, the PBN is used for steady-state and perturbation simulations to evaluate model behavior under disease and therapeutic contexts. Genetic mutations, knockouts, and drug treatments are modeled by fixing node states or assigning probabilistic activity levels that reflect partial efficacy. Differences in optimized rule probabilities and simulated phenotypes across conditions are examined to identify regulatory rewiring and pathway dependencies associated with specific mutation backgrounds or treatments.

The optimized model also enables predictive analysis of therapeutic response. Models trained on single-drug perturbation data can be used to simulate drug combinations and to predict relative phenotypic changes across genomic contexts. Model predictions are evaluated by comparison with independent experimental datasets and published evidence. In addition, simulated phenotype scores can be linked to clinical outcomes, such as overall survival, to assess the model’s ability to capture disease progression and treatment response.

### Datasets

#### The Beat AML dataset

To optimize and evaluate the augmented model for AML drug response prediction, the Beat AML dataset was used, which comprises a cumulative cohort of 805 AML patients (942 specimens) (32). WES/targeted sequencing mutation calls, inhibitor AUC values, and clinical summary were retrieved from https://biodev.github.io/BeatAML2/ in December 2025.

#### The FPMTB dataset

The functional precision medicine tumor board (FPMTB) dataset was used to externally validate the model, which contains *ex vivo* drug-response and multiomics profiling data for 252 samples from 186 patients with AML (33). Mutation data and drug response DSS scores were retrieved from https://zenodo.org/records/7370747 in December 2025.

#### Data processing

For both datasets, raw mutation calls and drug-response measurements were processed following the same procedures described in the original publications. Briefly, somatic mutation calls were used to define patient-specific mutation profiles. For drug-response, we focused on five targeted drugs for AML (FDA-approved or clinically investigated): Venetoclax, Entospletinib, Ibrutinib, Selumetinib, and Midostaurin — which represent key inhibitors in AML treatment strategies (reviewed in (34)). For both datasets, we created cohorts where patient response data was available for one of these treatments. Drug-response measurements (AUC for Beat AML and DSS for FPMTB) were reversed. and normalized using min–max scaling to the [0, 1] range, enabling direct comparison between experimental responses and simulated phenotypic outputs.

Patients were stratified into mutation-defined subgroups based on the presence or absence of recurrent AML drivers, specifically *FLT3*-ITD, *NPM1*, and *DNMT3A. NRAS* mutation status is also considered in the *NRAS*-extended model. These mutation profiles were used to define distinct patient groups for model training, simulation, and downstream analysis. They were mapped to model genes and encoded as fixed node states based on curated gene roles (1 for oncogene and 0 for tumor suppressor genes). Drug-response data were aligned to the corresponding inhibitors and samples, and cohorts were split according to the original Beat AML wave definitions (wave 1+2 for training; wave 3+4 for evaluation). Due to the limited sample size in the FPMTB data, we include only mutation groups with at least five patients to reduce variability in drug response.

Cross validation

## Author contributions statement

The work was initially proposed and conceptualized by GQ and JG. LL developed the workflow, conducted data analyses, and wrote the initial draft of the manuscript. YZ designed and implemented the knowledge graph query and model construction methods. BA contributed to the conceptualization and methodological design of the study, and implemented BN and PBN simulation methods. GQ provided supervision throughout the project. All authors reviewed, revised, and approved the final version of the manuscript.

## Acknowledgments

This project was supported by NIH/NCI grants U01CA282109, R01CA296872, R01CA270210 and 5P41EB023912. Support for this work was also provided by the National Center for Advancing Translational Sciences, National Institutes of Health, through the Biomedical Data Translator program, award 1OT2TR005706. The content is solely the responsibility of the authors and does not necessarily represent the official views of the NIH. We gratefully acknowledge the late Professor Ilya Shmulevich from the Institute for Systems Biology– a pioneer of probabilistic boolean networks, an esteemed mentor, and a dear friend for his inspiring guidance for this work.

