## Supplementary materials for "KGBN: Augmenting and optimizing logical gene regulatory networks using knowledge graphs"

### Supplementary Figures

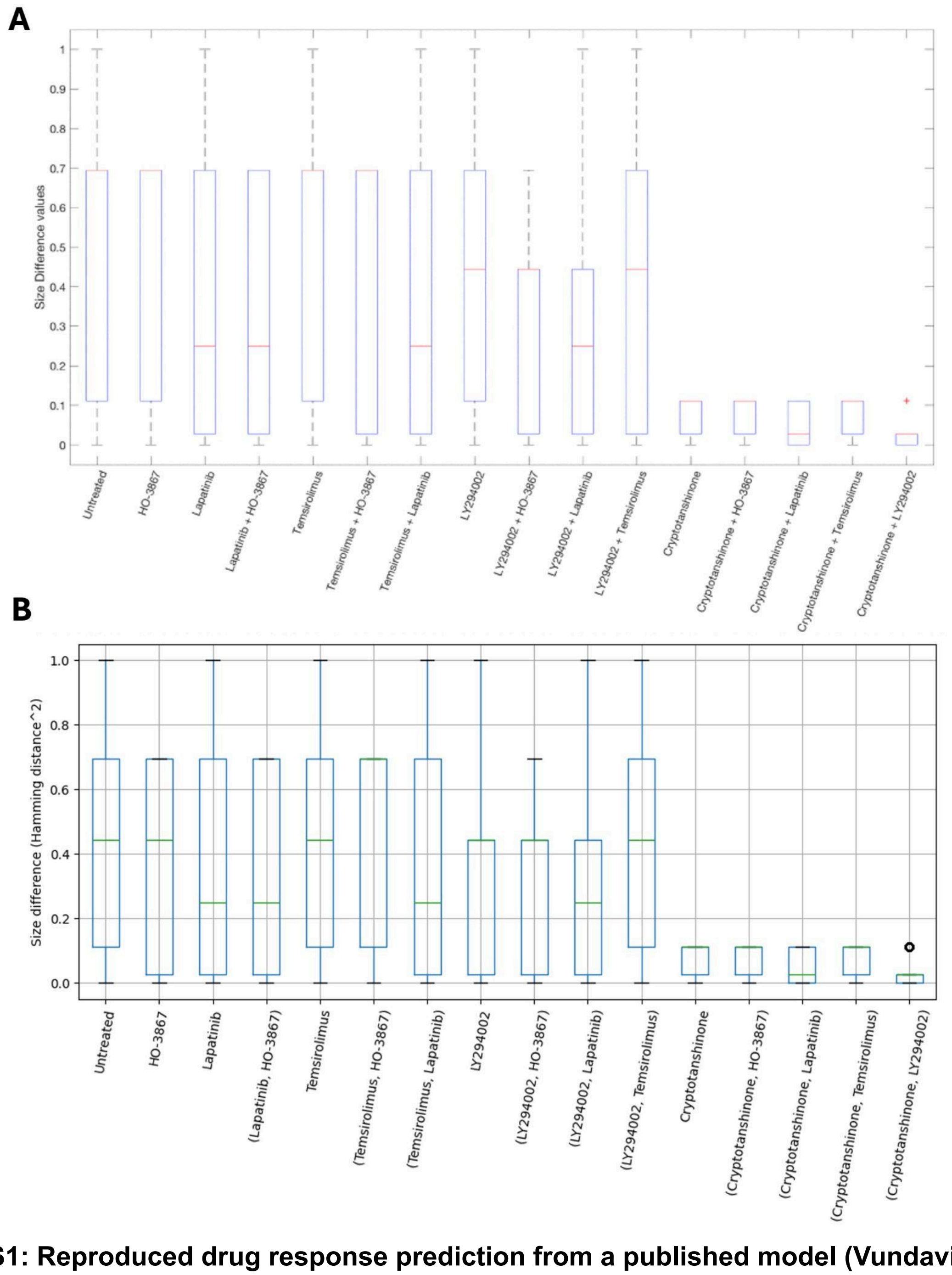


**Figure S1: Reproduced drug response prediction from a published model (Vundavilli et al. 2020).** (A) Fig. 5 from the original publication. (B) KGBN reproduced results.

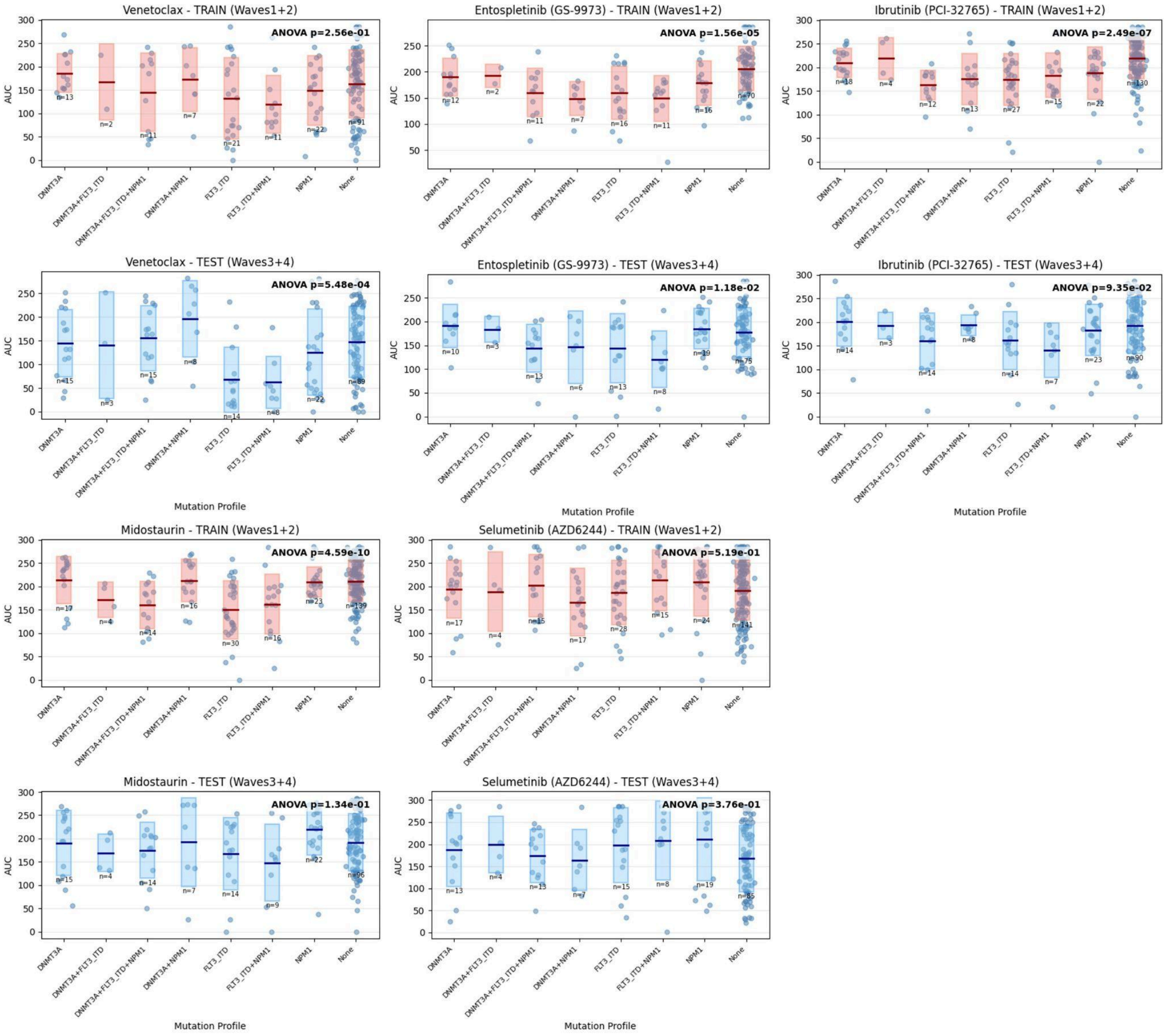


**Figure S2: Drug-response data used to optimize the base model.**

Averaged AUC of the 5 drugs for patients with similar mutational status of *FLT3*, *NPM1*, *DNMT3A*. Red: training set from Beat AML wave 1+2, blue: test set from Beat AML wave 3+4.

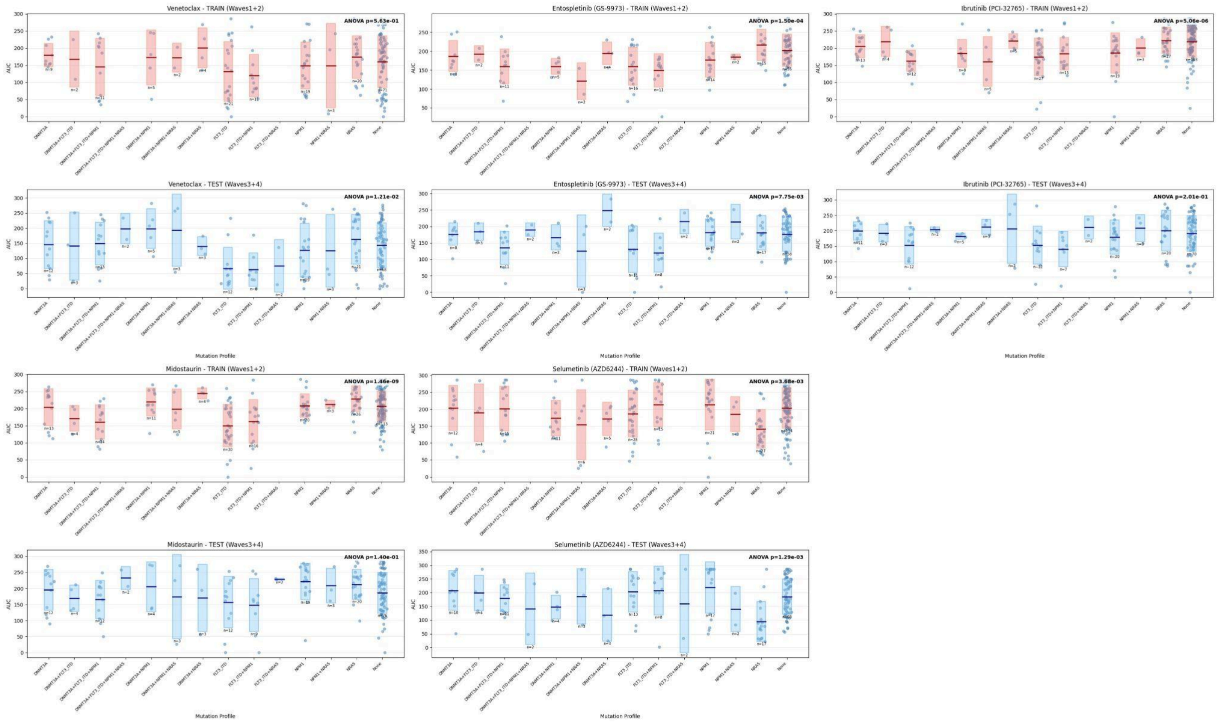


**Figure S3: Drug-response data used to optimize the *NRAS*-extended model.**
Averaged AUC of the 5 drugs for patients with similar mutational status of *FLT3*, *NPM1*, *DNMT3A,* and *NRAS*. Red: training set from Beat AML wave 1+2, blue: test set from Beat AML wave 3+4.

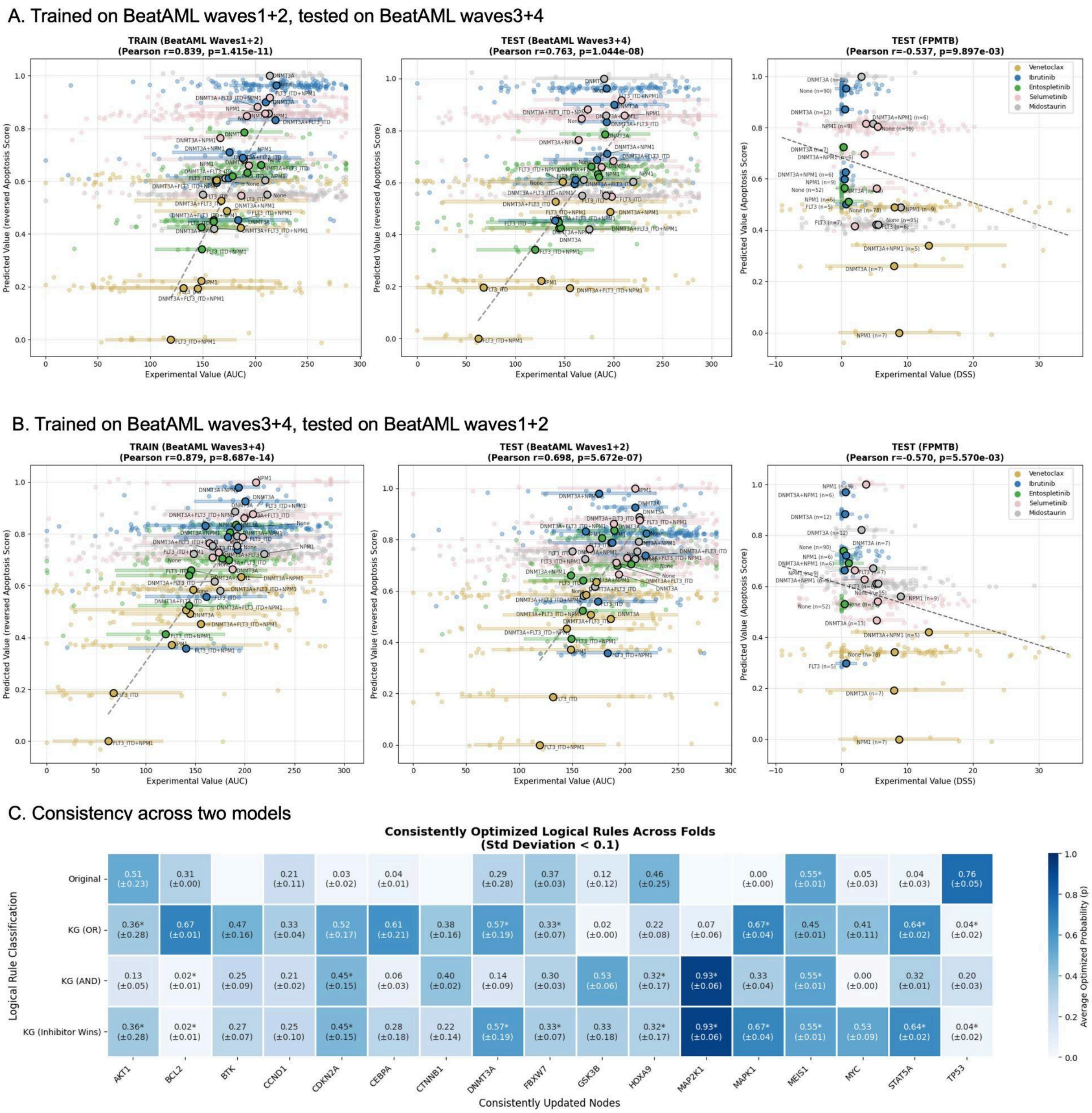


**Figure S4: Cross validation of the extended model on train-test splits.**
(A) Performance of the model (extended to drug targets) trained on BeatAML waves1+2 data, tested on BeatAML waves3+4 and external FPMTB data. (B) Performance of the model (extended to drug targets) trained on BeatAML waves3+4 data, tested on BeatAML waves1+2 and external FPMTB data. (C) Comparison of probabilities across models in (A) and (B).

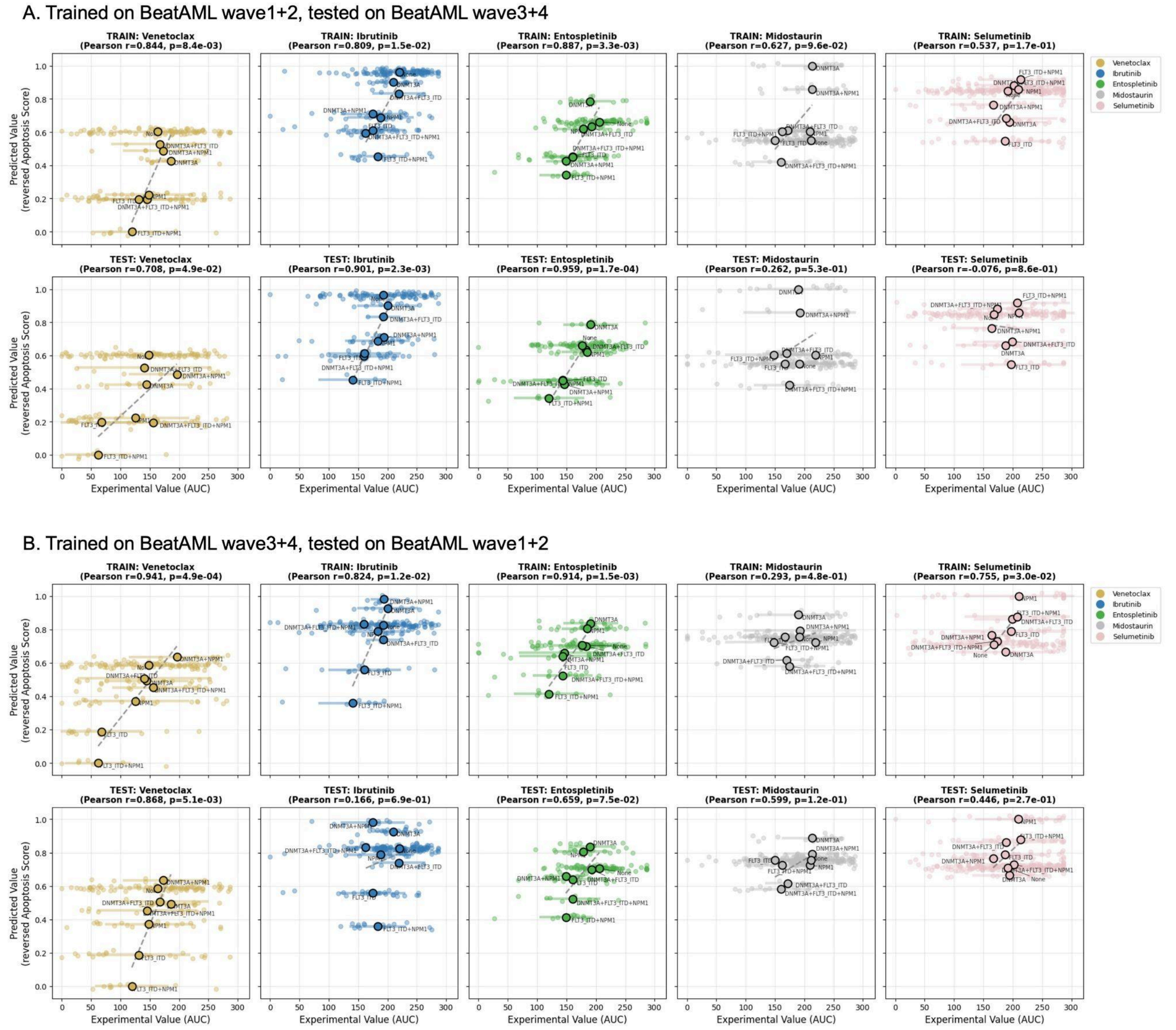


**Figure S5: Cross validation of the extended model on train-test splits, showing individual drugs.**

(A) Performance of the model (extended to drug targets) trained on BeatAML waves1+2 data, tested on BeatAML waves3+4 data. (B) Performance of the model (extended to drug targets) trained on BeatAML waves3+4 data, tested on BeatAML waves1+2 data. Both panels show results on individual drugs (colored differently.)

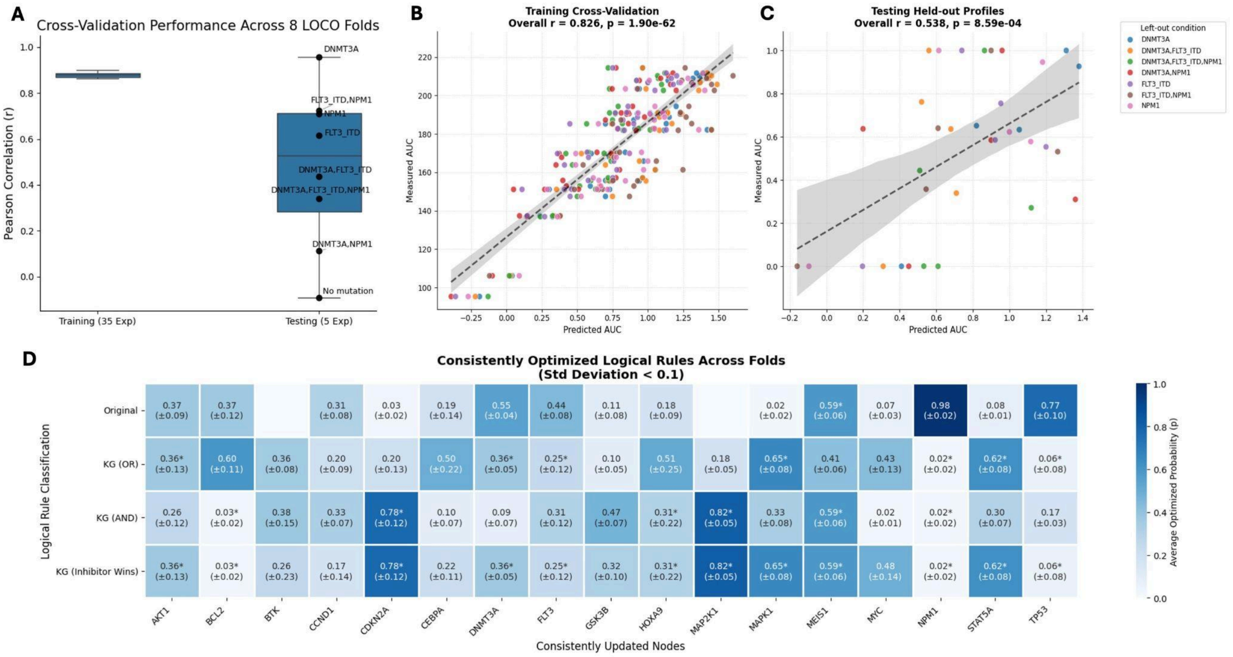


**Figure S6: Leave-one-mutation-out (LOCO) cross-validation of the model extended by drug targets.**

(A) Distribution of Pearson correlations across training and testing folds. (B) Aggregated predicted versus measured AUC for training folds. (C) Aggregated predicted versus measured AUC for the held-out mutation profiles. (D) Average optimized rule probabilities for rules with standard deviation below 0.1 across folds except no mutation.

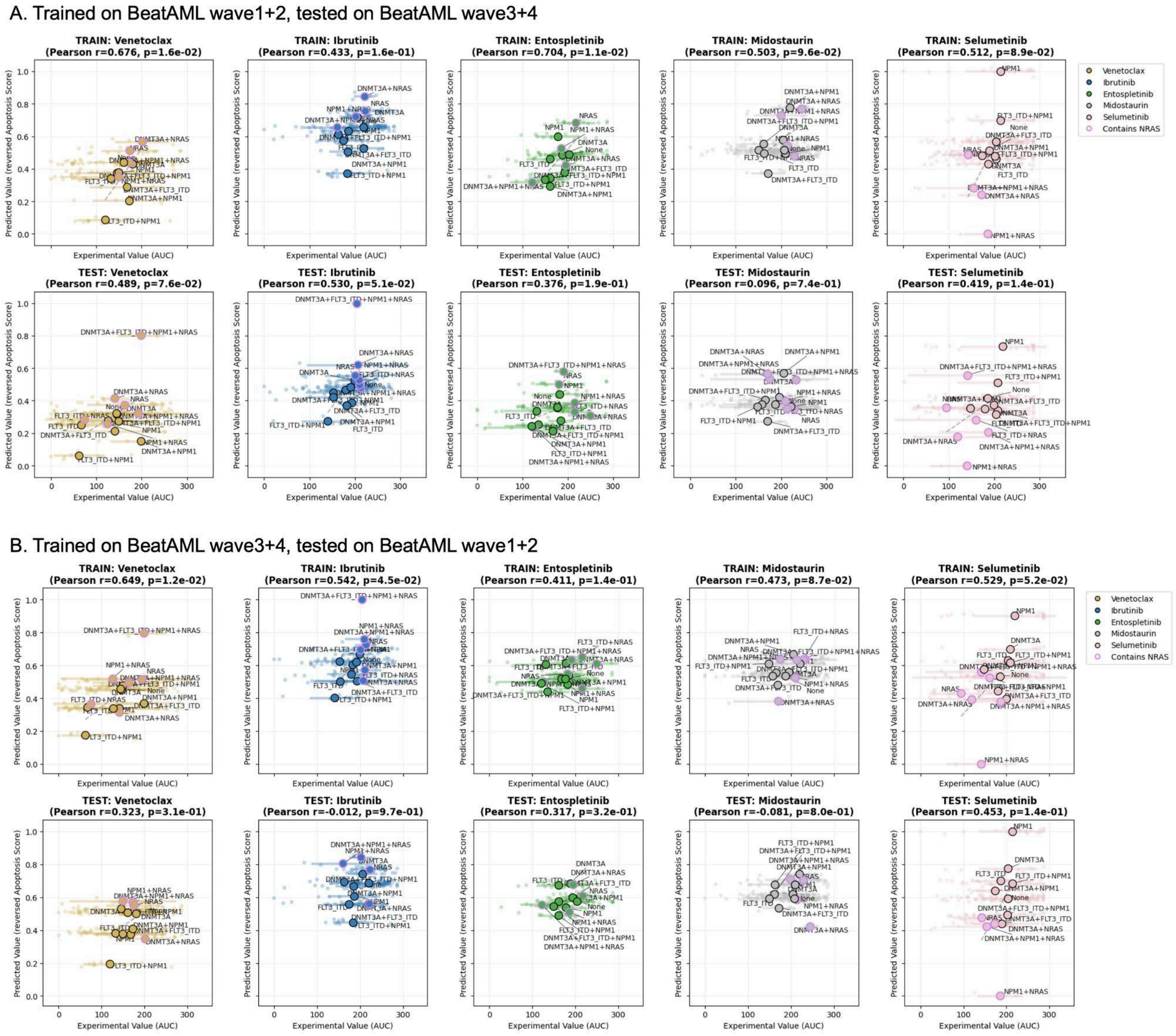


**Figure S7: Cross validation of the *NRAS*-extended model on train-test splits, showing individual drugs.**

(A) Performance of the model (extended to drug targets and *NRAS*) trained on BeatAML waves1+2 data, tested on BeatAML waves3+4 data. (B) Performance of the model (extended to drug targets and *NRAS*) trained on BeatAML waves3+4 data, tested on BeatAML waves1+2 data. Both panels show results on individual drugs (colored differently.)

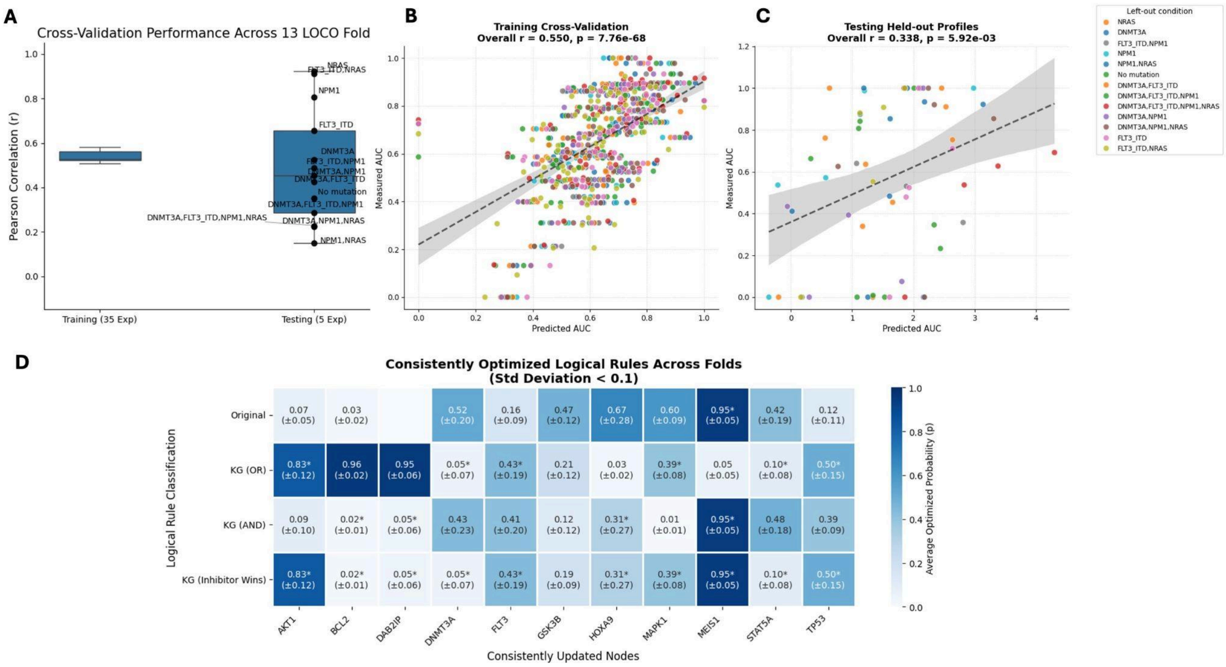


**Figure S8: LOCO cross-validation of the model extended by drug targets and *NRAS*.** (A) Distribution of Pearson correlations across training and testing folds. (B) Aggregated predicted versus measured AUC for training folds. (C) Aggregated predicted versus measured AUC for the held-out mutation profiles. (D) Average optimized rule probabilities for rules with standard deviation below 0.1 across folds except no mutation.

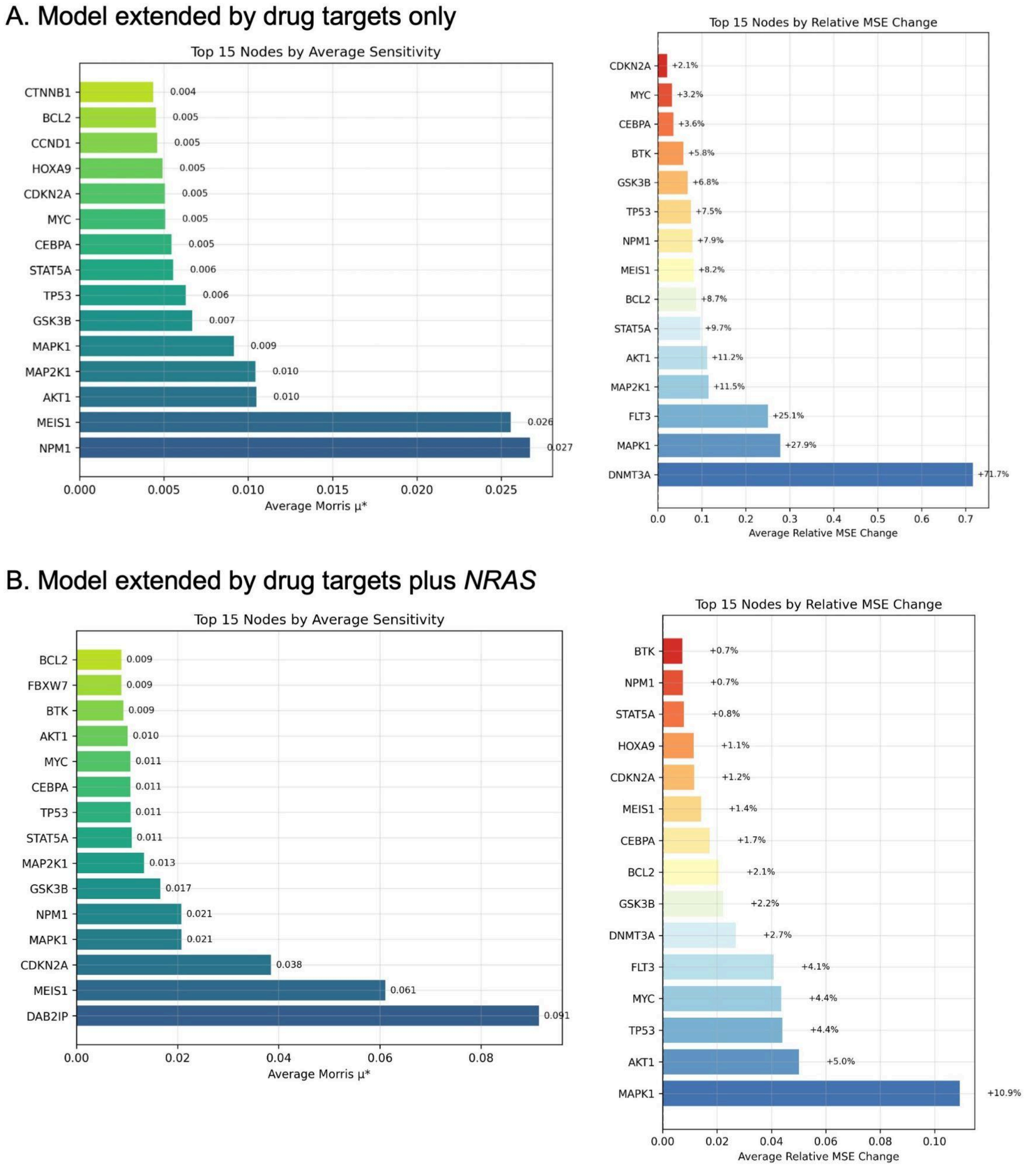


**Figure S9: Sensitivity analysis of the optimized model.**
Left panel: Global output sensitivity (Morris μ ), which quantifies the absolute effect that fluctuations in a node's rule probabilities have on the steady-state apoptosis prediction; Right panel: Model fit robustness (Relative MSE Change). For each multi-rule node, we perturbed its optimized probability distribution by 10% and measured the resulting degradation in predictive accuracy.

### Supplementary Tables

#### Table S1: Relations retrieved from SIGNOR knowledge graph.

| source | target | direction | predicate | Knowledge_Source | publications | score |
|---|---|---|---|---|---|---|
| AKT1 | AKT1 | activate | up-regulates activity | SIGNOR | 16549426 | 0.2 |
| AKT1 | BTK | inhibit | down-regulates quantity by destabilization | SIGNOR | 23754751 | 0.296 |
| AKT1 | DAB2IP | inhibit | down-regulates activity | SIGNOR | 27858941 | 0.497 |
| AKT1 | FBXW7 | activate | up-regulates activity | SIGNOR | 21620836 | 0.41 |
| AKT1 | GSK3B | inhibit | down-regulates activity | SIGNOR | 15829723 | 0.793 |
| AKT1 | MAPK1 | activate | up-regulates activity | SIGNOR | 20693286 | 0.532 |
| AKT1 | NPM1 | inhibit | down-regulates activity | SIGNOR | 25071014 | 0.534 |
| BTK | BTK | activate | up-regulates | SIGNOR | 8630736 | 0.2 |
| BTK | STAT5A | activate | up-regulates activity | SIGNOR | 11413148 | 0.468 |
| CEBPA | CEBPA | activate | up-regulates quantity | SIGNOR | 11283671 | 0.2 |
| CEBPA | MYC | inhibit | down-regulates quantity by repression | SIGNOR | 12032779 | 0.49 |
| CEBPA | SOX4 | inhibit | down-regulates | SIGNOR | 24183681 | 0.382 |
| DAB2IP | GSK3B | activate | up-regulates activity | SIGNOR | 20080667 | 0.312 |
| DAB2IP | NRAS | inhibit | down-regulates activity | SIGNOR | 27858941 | 0.493 |
| DNMT3A | CCND1 | inhibit | down-regulates quantity by repression | SIGNOR | 19786833 | 0.504 |
| DNMT3A | CDKN2A | inhibit | down-regulates quantity by repression | SIGNOR | 26350239 | 0.378 |
| DNMT3A | HOXA9 | inhibit | down-regulates quantity by repression | SIGNOR | 24280869 | 0.338 |
| DNMT3A | MEIS1 | inhibit | down-regulates quantity by repression | SIGNOR | 28288143 | 0.328 |
| FBXW7 | DAB2IP | inhibit | down-regulates quantity by destabilization | SIGNOR | 27858941 | 0.328 |
| FBXW7 | MYC | inhibit | down-regulates quantity | SIGNOR | 20852628 | 0.762 |
| FLT3 | AKT1 | activate | up-regulates activity | SIGNOR | 16266983 | 0.431 |
| FLT3 | CEBPA | inhibit | down-regulates quantity by repression | SIGNOR | 14592841 | 0.631 |
| FLT3 | FLT3 | activate | up-regulates activity | SIGNOR | 16627759 | 0.2 |
| FLT3 | MYC | activate | up-regulates quantity by expression | SIGNOR | 25280219 | 0.363 |
| FLT3 | STAT5A | activate | up-regulates activity | SIGNOR | 17356133 | 0.603 |
| GSK3B | BCL2 | inhibit | down-regulates quantity by destabilization | SIGNOR | 29107113 | 0.2 |
| GSK3B | CCND1 | inhibit | down-regulates | SIGNOR | 16504004 | 0.783 |
| GSK3B | CEBPA | activate | up-regulates activity | SIGNOR | 10567568 | 0.38 |
| GSK3B | GSK3B | activate | up-regulates activity | SIGNOR | 20331603 | 0.2 |
| GSK3B | MYC | inhibit | down-regulates quantity by destabilization | SIGNOR | 16023596 | 0.719 |
| GSK3B | TP53 | activate | up-regulates activity | SIGNOR | 11483158 | 0.727 |
| HOXA9 | MEIS1 | activate | up-regulates activity | SIGNOR | 9343407 | 0.62 |
| MAPK1 | BCL2 | activate | up-regulates quantity by stabilization | SIGNOR | 10669763 | 0.551 |
| MAPK1 | CEBPA | inhibit | down-regulates | SIGNOR | 14701740 | 0.352 |
| MAPK1 | ETV6 | inhibit | down-regulates | SIGNOR | 15060146 | 0.317 |
| MAPK1 | GSK3B | inhibit | down-regulates | SIGNOR | 16039586 | 0.384 |
| MAPK1 | MAPK1 | activate | up-regulates activity | SIGNOR | 1712480 | 0.2 |
| MAPK1 | MYC | activate | up-regulates activity | SIGNOR | 8386367 | 0.733 |
| MAPK1 | STAT5A | activate | up-regulates | SIGNOR | 10194762 | 0.759 |
| MAPK1 | TP53 | activate | up-regulates activity | SIGNOR | 15116093 | 0.777 |
| MYC | CCND1 | activate | up-regulates quantity by expression | SIGNOR | 12835716 | 0.495 |
| MYC | CDKN2A | inhibit | down-regulates quantity by repression | SIGNOR | 20551174 | 0.765 |
| MYC | DNMT3A | activate | up-regulates activity | SIGNOR | 19786833 | 0.711 |
| NPM1 | FBXW7 | activate | up-regulates quantity | SIGNOR | 18625840 | 0.483 |
| NPM1 | HOXA9 | inhibit | down-regulates quantity by repression | SIGNOR | 30205049 | 0.351 |
| STAT5A | DNMT3A | activate | up-regulates quantity | SIGNOR | 26059451 | 0.326 |
| SYK | BTK | activate | up-regulates activity | SIGNOR | 11226282 | 0.588 |
| SYK | FLT3 | activate | up-regulates activity | SIGNOR | 24525236 | 0.408 |
| SYK | SYK | activate | up-regulates activity | SIGNOR | 9820500 | 0.2 |
| TP53 | BCL2 | inhibit | down-regulates activity | SIGNOR | 19007744 | 0.748 |

**Table S2: Optimized PBNs.** Left: PBN extended to drug targets only; right: PBN extended to drug targets and *NRAS*; Underlined nodes and rules are from the original model.

| Node | PBN (extended to drug targets) | | PBN (extended to drug targets + *NRAS*) | |
|---|---|---|---|---|
| | **Rules** | **Probabilities** | **Rules** | **Probabilities** |
| **NRAS** | - | - | !DAB2IP | 1.000 |
| **DAB2IP** | - | - | !AKT1 & !FBXW7 | 0.900 |
| | - | - | !AKT1 \| !FBXW7 | 0.100 |
| **TP53** | CDKN2A | 0.806 | CDKN2A | 0.008 |
| | GSK3B & MAPK1 | 0.170 | GSK3B & MAPK1 | 0.958 |
| | GSK3B \| MAPK1 | 0.024 | GSK3B \| MAPK1 | 0.034 |
| **STAT5A** | BTK & FLT3 & MAPK1 | 0.331 | BTK & FLT3 & MAPK1 | 0.220 |
| | BTK \| FLT3 \| MAPK1 | 0.661 | BTK \| FLT3 \| MAPK1 | 0.012 |
| | FLT3 | 0.008 | FLT3 | 0.768 |
| **MAP2K1** | MAP2K1 & !MAPK1 | 0.865 | MAP2K1 & !MAPK1 | 0.126 |
| | MAP2K1 \| !MAPK1 | 0.135 | MAP2K1 \| !MAPK1 | 0.874 |
| **CEBPA** | !CTNNB1 & !FLT3 & !MAPK1 & ( CEBPA \| GSK3B \| MAP2K1 ) | 0.108 | !FLT3 & !MAPK1 & ( CEBPA \| GSK3B \| MAP2K1 ) | 0.015 |
| | !FLT3 | 0.038 | !FLT3 | 0.739 |
| | CEBPA & !CTNNB1 & !FLT3 & GSK3B & MAP2K1 & !MAPK1 | 0.031 | CEBPA & !FLT3 & GSK3B & MAP2K1 & !MAPK1 | 0.003 |
| | CEBPA \| !CTNNB1 \| !FLT3 \| GSK3B \| MAP2K1 \| !MAPK1 | 0.823 | CEBPA \| !FLT3 \| GSK3B \| MAP2K1 \| !MAPK1 | 0.243 |
| **KIT** | KIT | 1.000 | !DAB2IP & KIT | 0.843 |
| | - | - | !DAB2IP \| KIT | 0.157 |
| **MYC** | !CEBPA & !FBXW7 & !GSK3B & ( CTNNB1 \| FLT3 \| MAPK1 ) | 0.621 | !CEBPA & !FBXW7 & !GSK3B & ( FLT3 \| MAPK1 ) | 0.009 |
| | !CEBPA & CTNNB1 & !FBXW7 & FLT3 & !GSK3B & MAPK1 | 0.001 | !CEBPA & !FBXW7 & FLT3 & !GSK3B & MAPK1 | 0.031 |
| | !CEBPA \| CTNNB1 \| !FBXW7 \| FLT3 \| !GSK3B \| MAPK1 | 0.299 | !CEBPA \| !FBXW7 \| FLT3 \| !GSK3B \| MAPK1 | 0.640 |
| | MAPK1 & ( !FBXW7 \| !GSK3B ) | 0.079 | MAPK1 & ( !FBXW7 \| !GSK3B ) | 0.320 |
| **NPM1** | !AKT1 | 0.428 | !AKT1 | 0.993 |
| | NPM1 | 0.572 | NPM1 | 0.007 |
| **MAPK1** | AKT1 & MAP2K1 & MAP2K2 & MAPK1 | 0.292 | AKT1 & MAP2K1 & MAP2K2 & MAPK1 | 0.007 |
| | AKT1 \| MAP2K1 \| MAP2K2 \| MAPK1 | 0.706 | AKT1 \| MAP2K1 \| MAP2K2 \| MAPK1 | 0.496 |
| | FLT3 | 0.002 | FLT3 | 0.497 |
| **FLT3** | FLT3 & SYK | 0.118 | FLT3 & SYK | 0.598 |
| | FLT3 \| SYK | 0.332 | FLT3 \| SYK | 0.083 |
| | FLT3 | 0.550 | FLT3 | 0.319 |
| **AKT1** | AKT1 & FLT3 | 0.077 | AKT1 & FLT3 | 0.368 |
| | AKT1 \| FLT3 | 0.639 | AKT1 \| FLT3 | 0.193 |
| | FLT3 | 0.284 | FLT3 | 0.439 |
| **HOXA9** | !DNMT3A & !NPM1 | 0.141 | !DNMT3A & !NPM1 | 0.556 |
| | !DNMT3A \| !NPM1 | 0.146 | !DNMT3A \| !NPM1 | 0.011 |
| | !NPM1 | 0.713 | !NPM1 | 0.433 |
| **CDKN2A** | !DNMT3A & !MYC | 0.600 | !DNMT3A & !MYC | 0.672 |
| | !DNMT3A \| !MYC | 0.346 | !DNMT3A \| !MYC | 0.018 |
| | NPM1 | 0.054 | NPM1 | 0.311 |
| **DNMT3A** | DNMT3A | 0.567 | DNMT3A | 0.859 |
| | MYC & STAT5A | 0.047 | MYC & STAT5A | 0.067 |
| | MYC \| STAT5A | 0.386 | MYC \| STAT5A | 0.074 |
| **BTK** | !AKT1 & ( BTK \| SYK ) | 0.206 | !AKT1 & ( BTK \| SYK ) | 0.161 |
| | !AKT1 & BTK & SYK | 0.160 | !AKT1 & BTK & SYK | 0.001 |
| | !AKT1 \| BTK \| SYK | 0.634 | !AKT1 \| BTK \| SYK | 0.838 |
| **FBXW7** | AKT1 & NPM1 | 0.338 | AKT1 & NPM1 | 0.189 |
| | AKT1 \| NPM1 | 0.261 | AKT1 \| NPM1 | 0.350 |
| | NPM1 | 0.401 | NPM1 | 0.461 |
| **GSK3B** | !AKT1 & !MAPK1 & ( GSK3B \| MAP2K1 ) | 0.153 | !AKT1 & !MAPK1 & ( DAB2IP \| GSK3B \| MAP2K1 ) | 0.012 |
| | !AKT1 & GSK3B & MAP2K1 & !MAPK1 | 0.593 | !AKT1 & DAB2IP & GSK3B & MAP2K1 & !MAPK1 | 0.654 |
| | !AKT1 \| GSK3B \| MAP2K1 \| !MAPK1 | 0.013 | !AKT1 \| DAB2IP \| GSK3B \| MAP2K1 \| !MAPK1 | 0.007 |
| | !AKT1 | 0.241 | !AKT1 | 0.327 |
| **BCL2** | !GSK3B & MAPK1 & !TP53 | 0.015 | !GSK3B & MAPK1 & !TP53 | 0.004 |
| | !GSK3B \| MAPK1 \| !TP53 | 0.682 | !GSK3B \| MAPK1 \| !TP53 | 0.800 |
| | MAPK1 & !TP53 | 0.304 | MAPK1 & !TP53 | 0.196 |
| **MEIS1** | !DNMT3A & HOXA9 | 0.553 | !DNMT3A & HOXA9 | 0.640 |
| | !DNMT3A \| HOXA9 | 0.447 | !DNMT3A \| HOXA9 | 0.360 |
| **CCND1** | ! ( DNMT3A \| GSK3B ) | 0.100 | ! ( DNMT3A \| GSK3B ) | 0.177 |
| | !DNMT3A & !GSK3B & ( CTNNB1 \| MYC ) | 0.345 | !DNMT3A & !GSK3B & ( CTNNB1 \| MYC ) | - |
| | CTNNB1 & !DNMT3A & !GSK3B & MYC | 0.184 | !DNMT3A & !GSK3B & MYC | 0.268 |
| | CTNNB1 \| !DNMT3A \| !GSK3B \| MYC | 0.371 | !DNMT3A \| !GSK3B \| MYC | 0.555 |
| **MAP2K2** | MAP2K2 | 1.000 | MAP2K2 | 1.000 |
| **SYK** | SYK | 1.000 | SYK | 1.000 |
| **SOX4** | !CEBPA | 1.000 | !CEBPA | 1.000 |
| **ETV6** | !MAPK1 | 1.000 | !MAPK1 | 1.000 |

**Table S3: Cross validation of the extended model on leave-one-combination-out splits.** Each mutation combination was left out in training and tested on later.

A. Model extended by drug targets only

| mutation_profile | train_pearson_r | train_pearson_p | test_pearson_r | test_pearson_p | n_patients_min | n_patients_max |
|---|---|---|---|---|---|---|
| DNMT3A | 0.862 | 3E-11 | 0.957 | 1E-02 | 22 | 32 |
| FLT3_ITD,NPM1 | 0.834 | 5E-10 | 0.723 | 2E-01 | 19 | 25 |
| NPM1 | 0.881 | 3E-12 | 0.71 | 2E-01 | 35 | 45 |
| FLT3_ITD | 0.879 | 4E-12 | 0.616 | 3E-01 | 29 | 44 |
| DNMT3A,FLT3_ITD | 0.887 | 1E-12 | 0.437 | 5E-01 | 5 | 8 |
| DNMT3A,FLT3_ITD,NPM1 | 0.886 | 1E-12 | 0.341 | 6E-01 | 24 | 28 |
| DNMT3A,NPM1 | 0.899 | 2E-13 | 0.111 | 9E-01 | 13 | 24 |
| No mutation | 0.87 | 1E-11 | -0.093 | 9E-01 | 145 | 235 |

B. Model extended by drug targets plus *NRAS*

| mutation_profile | train_pearson_r | train_pearson_p | test_pearson_r | test_pearson_p | n_patients_min | n_patients_max |
|---|---|---|---|---|---|---|
| NRAS | 0.509 | 1E-05 | 0.923 | 3E-02 | 32 | 47 |
| FLT3_ITD,NRAS | 0.508 | 2E-05 | 0.911 | 3E-02 | 2 | 2 |
| NPM1 | 0.529 | 6E-06 | 0.806 | 1E-01 | 31 | 39 |
| FLT3_ITD | 0.522 | 8E-06 | 0.655 | 2E-01 | 27 | 42 |
| DNMT3A | 0.523 | 8E-06 | 0.526 | 4E-01 | 16 | 25 |
| FLT3_ITD,NPM1 | 0.528 | 6E-06 | 0.488 | 4E-01 | 19 | 25 |
| DNMT3A,NPM1 | 0.561 | 1E-06 | 0.453 | 4E-01 | 8 | 15 |
| DNMT3A,FLT3_ITD | 0.519 | 1E-05 | 0.425 | 5E-01 | 5 | 8 |
| No mutation | 0.636 | 1E-08 | 0.351 | 6E-01 | 113 | 189 |
| DNMT3A,FLT3_ITD,NPM1 | 0.553 | 2E-06 | 0.286 | 6E-01 | 22 | 26 |
| DNMT3A,NPM1,NRAS | 0.577 | 5E-07 | 0.226 | 7E-01 | 5 | 9 |
| DNMT3A,FLT3_ITD,NPM1,NRAS | 0.582 | 4E-07 | 0.225 | 7E-01 | 2 | 2 |
| NPM1,NRAS | 0.551 | 2E-06 | 0.149 | 8E-01 | 4 | 6 |
